# Adeno-associated viruses (AAVs) induce dose-dependent neonatal ventriculomegaly following intracerebroventricular administration

**DOI:** 10.64898/2025.12.30.697113

**Authors:** Luke L. Liu, Ania Noguera, Ryan Humphries, Blake Zhou, Ryann M. Fame

## Abstract

Cell-type-specific expression of synthetic/endogenous proteins or genetic sequences has significantly advanced our understanding of the central nervous system (CNS). Adeno-associated virus (AAV)-delivery to cerebrospinal fluid (CSF) mediates transfection of target cells to enable sustained delivery of secretory proteins into the CSF, offering promising avenues for both CNS therapy and mechanistic studies. However, despite the advantages afforded by AAV tropism-based cellular selectivity and transgene delivery, both preclinical studies and clinical trials report short- and long-term adverse effects, particularly immune activation. Especially relevant for CSF biology, CNS immune insults raise the risk of CSF dysregulation, including hydrocephalus. These risks may be exacerbated in pediatric populations, where ongoing CNS development, including immature meninges, choroid plexus (ChP), and skull structures, may further impair any ability to compensate for CSF dysregulation. To systematically address these risks and provide guidelines for minimizing CNS immune insults by AAVs, we test the dose-dependent effects of intracerebroventricular (ICV) injections of 3 AAV serotypes (AAV2/5, AAV2/4, and AAV.PHP.eB) to neonatal (P0.5) CD1 mouse pups. Histological analysis verified AAV2/5 tropism limited to ChP epithelial cells (CPECs), whereas AAV.PHP.eB transfected both CPECs and ependymal regions. By contrast, AAV2/4 shows limited transfection in the brain. Further, we find that ICV injections of all 3 AAV serotypes at the high dose (4x10^9^ genome copy GC/pup) induced ventriculomegaly by P7.5, while the low dose (1x10^9^ GC/pup) was well tolerated. Additionally, high-dose AAV2/5 tropism became more permissive, transfecting the ChP and also ependymal cells and some neurons. Longitudinal MRI of P0.5 pups with high-dose AAV2/5 ICV injections highlighted quick progression of ventriculomegaly. CSF ELISA analysis detected elevated pro-inflammatory cytokine CCL2 in AAV2/5 and AAV2/4 high-dose groups, indicating CNS inflammation. Moreover, decreased CSF TTR concentrations in high-dose groups suggest ChP dysfunction. We further revealed that earlier in utero high-dose ICV AAV2/5 injections at E13.5 induced even more severe ventriculomegaly and that adult animals were also susceptible to ventriculomegaly after high-dose AAV ICV delivery. Taken together, our data emphasize critical safety considerations of CSF-based AAV delivery, particularly during brain development. Further, these results call for an optimized dosage for perinatal ICV AAV applications.

**Highlights:**

- CSF-administration of AAVs results in unique cellular tropism in neonatal brains.
- Intracerebroventricular AAV delivery induces dose-dependent neonatal ventriculomegaly.
- Neonatal ventriculomegaly by AAV overdose develops quickly and progressively.
- AAV overdose ventriculomegaly correlates with ChP/CSF dyshomeostasis and inflammation.
- Adult and embryonic brains are also susceptible to dose-dependent AAV-induced ventriculomegaly.

## 1. Introduction

Adeno-associated virus (AAV)-based gene therapy has emerged as a promising treatment for previously incurable conditions. Since the first human trial in 1996 for cystic fibrosis (Flotte et al. 1996), AAV technology has achieved landmark success, notably with the approval of systemic AAV9 (Zolgensma^TM^) for spinal muscular atrophy type 1 in infants (Mendell et al. 2017). This success has driven further efforts to enhance AAV design, including optimizing payload expression, refining cell-specific tropism, and reducing immunogenicity. For CNS application, intrathecal or intraventricular delivery of AAVs has become more commonly tested (Flotte 2024; Zhou et al. 2024). However, despite these advances and the growing number of clinical trials, the efficacy and safety profiles of these engineered AAVs remain inadequately characterized in the context of the developing brain. This gap in knowledge requires thorough investigations before their widespread application in pediatric neurological diseases. In alignment with these clinical findings, results using AAVs in preclinical models are often difficult to reproduce and can be associated with non-specific phenotypes (Chakrabarty et al. 2013; Mathiesen et al. 2020).

Hydrocephalus, characterized by CSF dyshomeostasis and often ventricular enlargement, is a prevalent and serious condition in the pediatric population. Its diverse etiologies include intraventricular hemorrhage, genetic defects, infection, and traumatic brain injury (Kahle et al. 2024). Importantly, viral infections are established contributors to hydrocephalus pathogenesis (Paulson et al. 2020), likely through inflammatory mechanisms that disrupt cerebrospinal fluid (CSF) dynamics and impair drainage pathways (Karimy et al. 2020). While AAV-mediated gene therapy has emerged as a promising therapeutic strategy for neurological disorders, its inherent viral nature presents a potential risk. Viral particles may trigger neuroinflammatory responses or directly disrupt CSF homeostasis (Koyuncu et al. 2013), potentially leading to ventricular enlargement. This paradoxical risk—that a viral vector designed to treat a neurological condition could exacerbate one—is particularly salient in neonates, whose developing neural structures exhibit heightened vulnerability to insult. Given that standard interventions for hydrocephalus, e.g., shunt placement, choroid plexus (ChP) coagulation, endoscopic third ventriculostomy, remain highly invasive and carry significant failure rates, novel AAV-based therapies must be developed with stringent safety profiles and serotype-specific tropism evaluations to avoid iatrogenic harm in this susceptible population.

The fields of CSF AAV delivery coupled with pediatric gene therapy are therefore poised at a critical juncture, balancing the considerable promise of AAVs against the unique vulnerability of the developing brains to conditions such as hydrocephalus. To safely realize the potential of these therapies and to generate reliable preclinical models, a deeper understanding of AAV performance in a neonatal context is essential. To address this need, we designed this study to systematically characterize the effects of three preclinically relevant AAV serotypes following neonatal intracerebroventricular injections in mice. We anticipate that the findings from this investigation will improve the reliability of preclinical models by avoiding unintended effects of using AAV tools and provide essential preclinical data to guide the applications of safer and more effective AAV-based therapies for pediatric neurological disorders.

## 2. Materials and Methods

### 2.1. Mice

All animal procedures were performed in accordance with protocols approved by the Stanford University Administrative Panel on Laboratory Animal Care (APLAC) [Protocol #34330]. This work was conducted under Stanford University’s Animal Welfare Assurance #D16-00134 (A3213-01) within an AAALAC International accredited facility (Unit #000679). CD1 mice were obtained from Charles River Laboratories and housed under standard conditions with a 12-hour light/dark cycle (lights on at 7:00 a.m., off at 7:00 p.m.) and provided food and water ad libitum.

To obtain newborn pups on postnatal day 0 (P0), timed-pregnant CD1 female mice at embryonic day 17.5 (E17.5) were monitored during the evenings and early mornings to determine the exact timing of parturition. Newborn pups were subjected to experimental procedures within three hours of birth verification. In a separate cohort, timed-pregnant CD1 females at E12.5 were acclimated for 24 hours prior to any procedures. The morning after a vaginal plug was observed is denoted at day E0.5.

### 2.2. Neonatal intracerebroventricular injections

Anesthesia was induced in postnatal day 0.5 (P0.5) pups by placement on a pre-cooled aluminum plate over ice for approximately three minutes. Upon the absence of a response to tactile stimulation, the pup was transferred to a pre-cooled Cunningham adaptor equipped with ear bars for stabilization. The head was positioned to align the sagittal suture (between bregma and lambda) parallel to the surface of the adaptor.

A Hamilton syringe, mounted on a stereotaxic arm, was first positioned and zeroed at the midpoint of the bregma-lambda suture. The needle was first moved laterally by 1.0-1.1 mm. This distance corresponds to the point where a horizontal line from the bregma-lambda midpoint intersects a line drawn from lambda to the eye (**Fig. 1A**). This intersection defined the skull surface coordinate for needle penetration, which was used for injections in CD1 pups up to P0.5. To account for postnatal head growth, the lateral coordinate was adjusted for older pups: 1.1-1.2 mm for P1.5, 1.2-1.3 mm for P2.5, and 1.3-1.4 mm for P3.5. Once the skull surface coordinate was determined, the syringe was lowered to superficially penetrate the skin and skull, establishing the injection trajectory before the final ventricular delivery. The syringe was then advanced to a depth of 2.5 mm to access the lateral ventricle. A total of 1 µL of AAV solution was infused into the CSF at a rate of 1 µL/min (information regarding AAVs used in this study was provided in Table 1). The needle remained unmoved for 3 minutes to permit sufficient diffusion and prevent backflow. This method allows for an accurate delivery to the neonatal brain ventricles (Fig.1B). Two different viral concentrations were tested for each of the three AAV serotypes: 1 × 10¹² GC/mL (low dose), and 4 × 10¹² GC/mL (high dose). Following infusion, the needle was left in place for an additional three minutes to minimize backflow and was subsequently withdrawn slowly. Post-operative pups were recovered on a warming pad for three minutes before being returned to the home cage. Post-operative analgesia was provided via subcutaneous injections of buprenorphine in polymer for slow release, in strict adherence to the IACUC-approved protocol.

**Figure 1.**
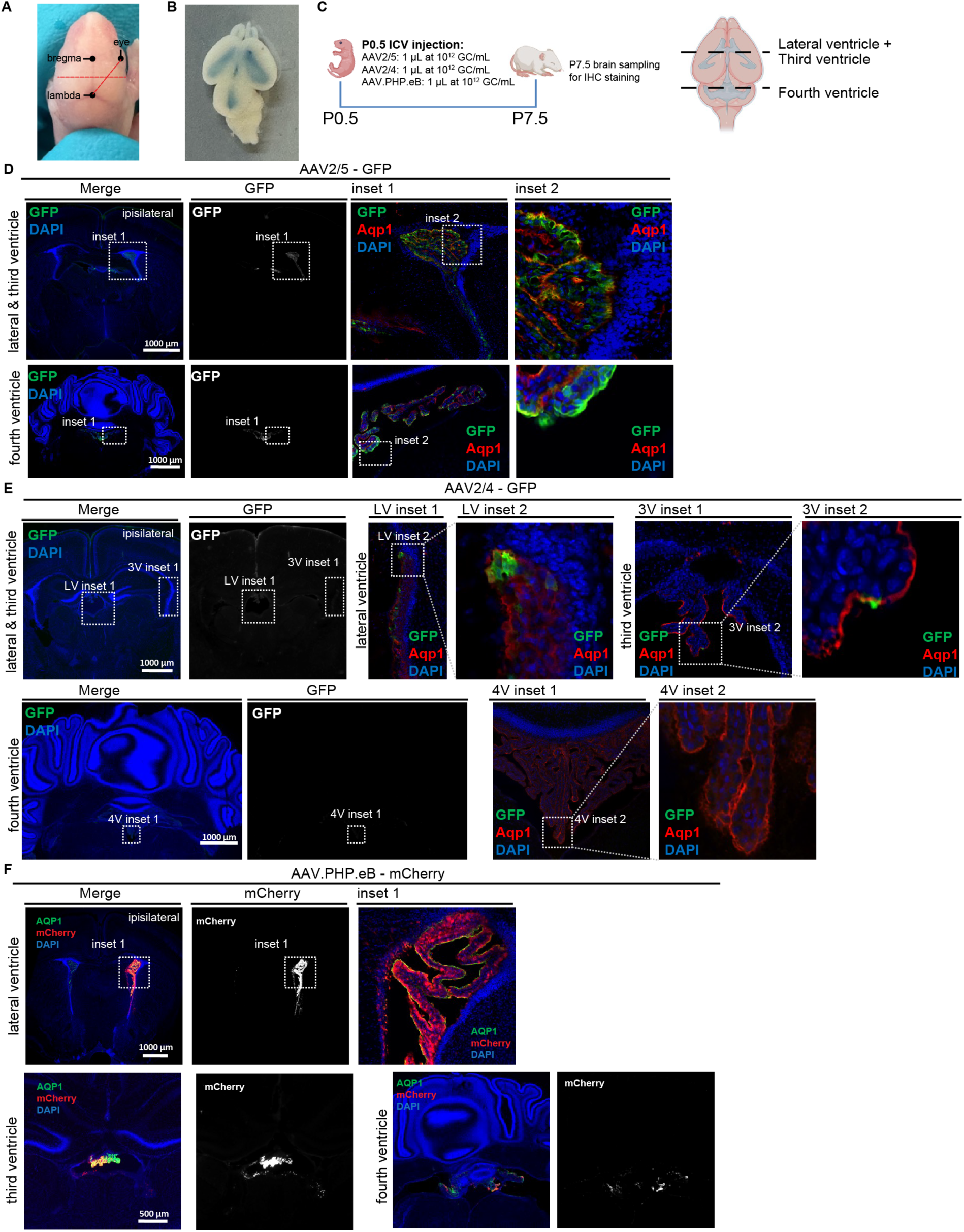
AAV2/5, AAV2/4, and AAV.PHP.eB exhibit shared and differential cellular tropism following neonatal ICV injection. **(A)** Dorsal view of a P0.5 pup head, showing skull landmarks used for reproducible and accurate intracerebroventricular (ICV) injection. The injection site (XY coordinates) is marked by the intersection of the two dashed lines: one horizontal line at the level of bregma/lambda midpoint and one line connecting the lambda and the eye. **(B)** Dorsal view of a P0.5 pup brain immediately after ICV injection of Evans Blue dye, confirming successful ventricular delivery. **(C)** Experimental timeline for AAV injection at P0.5 and analysis at P7.5. **(D)** Transfection pattern of AAV2/5 (GFP) in a P7.5 brain following ICV injection at P0.5. Insets show progressively higher-magnification views of transgene expression in the lateral/third ventricles and fourth ventricle, respectively. **(E)** Transfection pattern of AAV2/4 (GFP) in a P7.5 brain. Insets show progressively higher-magnification views of the lateral/third and fourth ventricles, respectively. Transfection pattern of AAV.PHP.eB (mCherry) in a P7.5 brain. Insets show progressively higher-magnification views of the lateral and fourth ventricles, respectively.

**Table 1.** Information regarding AAVs used in this study.

| AAV structures | Identifier |
| --- | --- |
| AAV2/5-CMV-GFP | SignaGen Laboratories SL100819 |
| AAV2/4-CMV-GFP | SignaGen Laboratories SL101269 |
| AAV.PHP.eB-CMV-mCherry | SignaGen Laboratories SL116133 |

To model the distribution of AAV particles based on their biophysical properties—which include a diameter of approximately 26 nm and a negative surface charge (Wright 2008)—we performed control injections with 1 µL fluorescent FluoSpheres carboxylate-modified microspheres (20 nm diameter, negative surface charges; Invitrogen, F8787). These microspheres were injected into P0.5 pups using the identical ICV injection procedure described above. Brains were extracted 24 hours post-injection for histological analysis. Post-operative analgesia was provided via subcutaneous injections of buprenorphine in polymer for slow release, in strict adherence to the IACUC-approved protocol.

### 2.3. In utero intracerebroventricular injections

Timed-pregnant CD1 mice at embryonic day 13.5 (E13.5) were anesthetized with 2.5% isoflurane and placed on a sterile, warm surgical pad. Under aseptic conditions, a laparotomy was performed to exteriorize the uterine horns. Individual embryos were gently stabilized to visualize the lateral ventricles through the uterine wall. A total volume of 1 µL of AAV2/5 solution containing 0.1% Evans blue was injected into the lateral ventricle of each targeted embryo using a beveled glass capillary needle. Two viral concentrations were tested: 1 × 10¹² GC/mL (low) and 4 × 10¹² GC/mL (high). Successful injection was confirmed by immediate visual observation of the bluish hue within the ventricular space, indicating proper delivery of the viral suspension. Following the injections, the uterine horns were carefully returned to the abdominal cavity. The abdominal muscle layer and skin incision were sutured separately. Dams were recovered from anesthesia on a warm pad and monitored post-operatively. Post-operative analgesia was provided via subcutaneous injections of buprenorphine in polymer for slow release, in strict adherence to the IACUC-approved protocol.

### 2.4. Adult intracerebroventricular injections

Adult CD1 mice were anesthetized with 5% isoflurane in the induction chamber and transferred onto a stereotaxic frame for head stabilization with a bite bar and ear bars. Following eye ointment application and fur removal, a midline cut was made on the scalp to reveal the bregma, which represents the reference coordinate of the skull. A cranial burr hole was then drilled on the right side 0.8 mm lateral of the bregma. A Hamilton syringe prefilled with AAV2/5 solution at 5x10^12^ GC/mL was lowered down by 2.5 mm through the burr hole to reach the lateral ventricle. The AAV solution (1 µL as low dose, 4 µL as high dose) was slowly injected to the CSF, and the needle remain unmoved in the ventricle for an additional 5 min before withdrawal. Upon completion of injections, the incision was closed by suturing followed by topical antibiotic ointment applications. ICV injection of FluoSpheres (1 µL) used the same procedure. Post-operative analgesia was provided via subcutaneous injections of buprenorphine in polymer for slow release, in strict adherence to the IACUC-approved protocol.

### 2.5. Histology and Microscopy

Dissected brain samples were fixed in 4% paraformaldehyde (PFA) in phosphate-buffered saline (PBS) for 24 hours at 4°C. Following fixation, tissues were washed in PBS and cryoprotected by immersion in 30% sucrose solution for 48 hours. Samples were then incubated overnight in a 1:1 mixture of 30% sucrose and Neg-50™ frozen section medium before being embedded in Neg-50™, equilibrated for one hour, and rapidly frozen. Frozen blocks were stored in sealed bags at -80°C until sectioning.

Coronal sections (40 µm thickness) were obtained using a Leica CM1860 cryostat. For immunohistochemistry (IHC), free-floating sections were blocked/permeabilized in blocking solution (PBST with 5% normal goat/donkey serum) supplemented with 0.3% Triton X-100. Sections were then incubated overnight at 4°C with primary antibodies diluted in blocking solution. After three washes in PBST, sections were incubated for 2 hours at room temperature with species-appropriate secondary antibodies conjugated to fluorescent dyes, diluted in blocking solution. Following counterstaining with Hoechst 33342, sections were mounted with Fluoromount-G™ aqueous mounting medium. Antibody details are provided in **Table 2**.

**Table 2.** Information regarding antibodies used in this study.

| Target | Host species | Identifier | Dilution |
| --- | --- | --- | --- |
| GFP | Rabbit | Invitrogen: A-6455 | 1:1000 |
| mCherry | Chicken | Aves Labs: MCHERRY-0020 | 1:1000 |
| AQP1 | Rabbit | Chemicon: AB2219 | 1:1000 |
| AQP1 | Mouse | Santa Cruz Biotechnology: sc-32737 | 1:500 |
| TTR | Rabbit | Invitrogen: MA5-47192 | 1:2000 |
| Iba-1 | Rabbit | FUJIFILM Wako: 019-19741 | 1:500 |
| Vinculin | Rabbit | Cell Signaling Technology: 13901S | 1:2000 |

For cell density quantification, brain sections were first imaged at low magnification (2x, Keyence BZ-X800) to identify regions of interest (ROIs). High-resolution z-stacks were then acquired within these ROIs using a confocal microscope (20x, Zeiss LSM T-PMT). Confocal images were analyzed in FIJI. ROIs were delineated using the freehand tool, and the corresponding areas were measured. Cells immunopositive for specific markers (e.g., Iba1) were manually counted within each ROI using the Cell Counter plugin. Cell density was calculated as the total number of cells divided by the area of the ROI.

### 2.6. Quantification of FluoSphere uptake in brain tissues

To quantify FluoSphere uptake, brain tissues were collected from P1.5 mouse pups or adult mice 24 hours after intracerebroventricular (ICV) injections. Anesthetized with a ketamine/xylazine solution and transcardially perfused with ice-cold phosphate-buffered saline (PBS), the ChP, ependyma, cortex, and meninges were then microdissected. Collected tissues were homogenized in a gentle lysis buffer (1% Triton X-100, 50 mM Tris-HCl, 150 mM NaCl, 1X Halt protease inhibitor cocktail, pH 7.4) using tissue grinders. Homogenates were incubated overnight at 4°C on a rotating shaker to ensure complete lysis.

Following the incubation, the lysates were clarified, and aliquots were taken for both fluorescence measurement and total protein quantification. Fluorescence intensity was measured using a Tecan Spark multimode microplate reader. A standard curve was generated from a serial dilution of FluoSpheres. Both standards and tissue lysate aliquots (5 µL) were loaded into a black 384-well plate. Fluorescence was measured with excitation and emission wavelengths set to 450 nm and 550 nm, respectively. The instrument’s Z-position and gain were optimized using the wells containing the highest standard concentration; 20 flashes per well were used for signal acquisition. FluoSphere uptake in each tissue was expressed as relative fluorescence intensity normalized to the total protein concentration, as determined by the BCA assay.

### 2.7. Magnetic resonance imaging (MRI)

Longitudinal in vivo MRI was performed on the same mouse pups at 2, 5, and 7 days following ICV injections performed at P0.5. Imaging was conducted on a 7.0-Tesla Bruker BioSpec 70/40 small-animal scanner (Bruker Corp., MA) operating at 300 MHz, equipped with a BGA-12S gradient insert (660 mT/m, 4570 T/m/s) and running ParaVision 360 software (version 3.6). Signal reception was achieved using a 4-element receive-only mouse CryoProbe™ paired with an 86 mm volume transmit coil. During imaging, pups were anesthetized with 3% isoflurane for induction and maintained on 1–1.5% isoflurane. Core body temperature was maintained at 37°C using a warm water-circulating pad with a feedback controller. Respiration was monitored throughout the procedure via a pneumatic pad (SA Instruments, NY) placed beneath the animal. Three-dimensional T2-weighted volumetric images were acquired using a RARE sequence with a Variable Flip Angle (VFL) technique. Key acquisition parameters were listed as follows: TR/TE = 1800/90.26 ms, FOV = 15 × 15 × 6.5 mm³, matrix = 150 × 150 × 65 (yielding 100 µm isotropic resolution), RARE factor = 43, NEX = 1.2, and a total scan time of 9 minutes 9 seconds. For volumetric analysis, the ventricles were manually segmented on every slice in the anterior-to-posterior direction using the open-source Horos software. This segmentation enabled three-dimensional reconstruction and precise quantification of ventricular volume.

### 2.8. CSF collection and ELISA

CSF was collected from P7.5 pups. Pups were fully anesthetized via intraperitoneal injection of a ketamine/xylazine solution and secured in a prone position on a customized Styrofoam platform using tape, orienting the head to present the cisterna magna. The neck area was placed under a dissection microscope. The skin and the three overlying muscle layers were carefully dissected away to expose the cisterna magna. Bleeding during the dissection was controlled using cotton swabs. Upon removal of the final muscle layer, the white-colored dura mater and the triangular cisterna magna were visualized.

A glass capillary needle, mounted on a micromanipulator for precise control, was carefully advanced into the cisterna magna. Gentle negative pressure was applied to draw CSF into the capillary. Collected CSF samples were expelled into low-binding microcentrifuge tubes and immediately centrifuged at 1,000 × g for 5 min at 4°C to pellet any cellular debris without inducing cellular lysis.

Samples with visible blood contamination were discarded. The resulting clarified CSF supernatants were stored at -80°C until analysis.

The concentrations of C-C motif chemokine ligand 2 (CCL2) and transthyretin (TTR) in the CSF were quantified using commercial ELISA kits (mouse CCL2 ELISA kit, BSKM1011, Bioss; mouse TTR ELISA kit, ab282297, Abcam), according to the manufacturers’ protocols. For CCL2 quantification, 5 µL of undiluted CSF was used for the initial incubation in antibody-coated wells. For TTR quantification, 1 µL of CSF was diluted 100-fold in the provided sample diluent prior to incubation with the antibody cocktail.

After completing all assay steps, the final absorbance was read at 450 nm. Sample concentrations were interpolated from the standard curve using a four-parameter logistic (4PL) curve fit in GraphPad Prism software.

### 2.9. Immunoblotting

Under ketamine/xylazine anesthesia, ChP tissues were isolated from P7.5 pups that received ICV injections AAV2/5-GFP on P0.5. ChP tissues were homogenized in RIPA buffer supplemented with 1X Halt protease inhibitor cocktail, and total protein concentrations were determined using a BCA assay.

Each sample containing 2 µg protein supplemented with 4X NuPAGE LDS Sample Buffer (Invitrogen, NP0007) and 2.5% dithiothreitol (DTT) were denatured at 95°C for 10 minutes.

Denatured ChP samples were loaded on NuPAGE 4–12% Bis-Tris Mini Protein Gels (Invitrogen, NP0329BOX) for electrophoresis with NuPAGE MES SDS Running Buffer (Invitrogen, NP0002) at 160V for 90 minutes. The proteins were subsequently transferred to a methanol-activated PVDF membrane using NuPAGE Transfer Buffer (Invitrogen, NP00061) at 30V for 1 hour. The membrane was blocked for 1 hour at room temperature with 5% non-fat dry milk in Tris-buffered saline with 0.1% Tween-20 (TBST). It was then incubated overnight at 4°C with primary antibodies against TTR and Vinculin diluted in TBST containing 5% bovine serum albumin (BSA). Following three 5-minute washes with TBST, the membrane was incubated for 2 hours at room temperature with an HRP-conjugated goat anti-rabbit secondary antibody in TBST with 5% BSA. After three additional TBST washes, the membrane was incubated with Amersham ECL Western Blotting Detection Reagents (Cytiva, DE) for 5 minutes and imaged with a GelDoc XR+ system (Bio-Rad, CA). To probe for GFP, the same membrane was incubated in Restore Western Blot Stripping Buffer (21059, Thermo Scientific, MA) for 30 minutes at 37°C. Following three TBST washes, the immunoblotting procedure (blocking, primary/secondary antibody incubation, ECL signal development, and imaging) was repeated as described.

Band intensities for TTR, GFP, and Vinculin were quantified using FIJI software and relative expressions of TTR and GFP were quantified by normalizing to Vinculin levels. Furthermore, relative TTR and GFP levels of individual ChP samples was plotted and analyzed by simple linear regression using GraphPad Prism 9. Full-membrane immunoblotting images were provided in **Fig. S5A-C**.

### 2.10. Minor inflammation and CSF metal analysis

To model sickness-like mild inflammation, LPS was administered to adult mice once via intraperitoneal injections at 50 µg/kg b.w. as previously described (Osterhout et al. 2022). Two hours after LPS dosing, mice were anesthetized with ketamine/xylazine solution for CSF collections (as described above). CSF samples were analyzed with inductively coupled plasma mass spectrometry (iCAP RQ, Thermo Scientific, MA). Briefly, 4 µL CSF was mixed into 5 mL matrix solution (0.1% HNO3 solution) to fit in the standard curves, which were generated from standard solutions containing K⁺ at 0.00, 1.20, 2.40, 4.80, 7.20, and 12.00 µM. Quality control was performed using R^2^ of standard curves (R^2^ of this batch was 0.9999).

### 2.11. Alcian Blue Staining

The expression of anionic glycans/proteoglycans, important brain extracellular matrix components (Schnaar et al. 2014), was evaluated using the Alcian Blue (pH 2.5) Stain Kit (H-3501, Vector Labs, CA). Cryosections of neonatal brains (20 µm thickness) were first hydrated in distilled water. Tissue sections were then incubated with 3% acetic acid solution for 3 minutes, followed by Alcian blue solution for 20 minutes at 37°C. After staining, sections were rinsed three times with acetic acid solution and then three times with distilled water (2 minutes per wash). A graded ethanol series (50%, 75%, 90%, 95%, and 100%; 5 minutes each) was used for dehydration. Finally, dehydrated sections were mounted with mounting medium and coverslips and imaged under brightfield using Keyence BZ-X800 microscope.

### 2.12. Statistics

Data are presented as mean ± standard deviation (SD). Statistical analyses were performed using GraphPad Prism software. Comparisons between two groups were analyzed using an unpaired, two-tailed Student’s t-test. For comparisons among more than two groups, one-way analysis of variance (ANOVA) was conducted, followed by Tukey’s post hoc test for multiple comparisons. A *p*-value of less than 0.05 was considered statistically significant. Significance levels are denoted as follows: \**p* < 0.05, \*\**p* < 0.01, \*\*\**p* < 0.001, and \*\*\*\**p* < 0.0001.

## 3. Results

### 3.1. AAV2/5, AAV2/4, and AAV.PHP.eB exhibit overlapping, but distinct cellular tropism in neonatal brains

To evaluate the cellular tropism of three AAV serotypes with ChP tropism in the neonatal brain, we performed intracerebroventricular (ICV) injections of AAV2/5, AAV2/4, or AAV.PHP.eB into newborn (P0.5) pups at a standard dose of 1×10⁹ genome copies (GC) (**Fig. 1A-C**). All vectors encoded a fluorescent reporter (**Fig. 1C**), and brains were analyzed one week post-injection (P7.5).

AAV2/5 exhibited highly specific tropism for choroid plexus epithelial cells (CPECs) across all ventricles (lateral, third, and fourth) (**Fig. 1D**). Specificity was confirmed by GFP signal co-localization with AQP1-positive cells. Notably, GFP intensity was markedly stronger in the ipsilateral lateral ventricle (LV) ChP compared to the contralateral side. In contrast, AAV2/4—a serotype previously reported to have ependymal specificity (Liu et al. 2005)—showed surprisingly low transfection efficiency in the neonatal brain in our hands (**Fig. 1E**). Transfected cells were primarily a small subset of CPECs in the ipsilateral LV, with only minimal signal in the contralateral LV, third ventricle (3V), fourth ventricle (4V), or ependyma. Additional signal was observed in the meninges, but not in ependyma (**Fig. 1E**).

AAV.PHP.eB, a serotype engineered for blood-brain-barrier crossing and broad parenchymal transduction (Chan et al. 2017), demonstrated robust tropism for CPECs following ICV injection (**Fig. 1F**). It also transfected ependymal cells and cells slightly deeper in the parenchyma along the ventricles.

Similar to AAV2/5, AAV2/4 transduction was largely ipsilateral. We did not observe the widespread cortical and hippocampal transduction previously reported in older mice of different strains (Mathiesen et al. 2020).

Collectively, these results reveal a shared primary tropism for CPECs across all three AAV serotypes after ICV injection in the neonatal brain. This common targeting, combined with the physical properties of viral particles and the extensive apical surface area of CPECs that contact CSF, suggests a unique capacity of the ChP to capture AAVs from the CSF. To test this capture mechanism directly, we administered FluoSpheres—biologically inert, nano-scale beads matching the approximate size and surface charge of AAVs—into the neonatal CSF (**Fig. S1A**). Histologically, FluoSpheres were abundantly captured by ipsilateral ChP tissue, with only sparse distribution in other regions such as the ependyma (**Fig. S1B**). Quantitative analysis confirmed the ChP as the dominant site of FluoSphere uptake in the neonatal brain (**Fig. S1C**). This preferential capture was even more pronounced in adult brains, where the ChP remained the dominant accumulation site (**Fig. S1D-1F**).

Together, these findings demonstrate serotype-specific cellular tropism in the neonatal brain, that is enriched on the ipsilateral side relative to the injection site, and a robust, inherent capacity of the ChP tissues to capture viral and viral-sized particles from the CSF.

### 3.2. AAVs induce dose-dependent neonatal ventriculomegaly

The continuous buildup of extracellular matrix (ECM) is fundamental to brain development and the establishment of its mechanical properties (Amin and Borrell 2020; Schnaar et al. 2014). For the brain ventricular system, a specific subset of ECM components—anionic glycans and their proteoglycan conjugates—is particularly critical; loss of these molecules lead to severe ventriculomegaly in animal models (Alonso et al. 1998; Angata et al. 2007; Weinhold et al. 2005). Using Alcian blue to visualize these anionic glycans, we observed their progressive accumulation in the neonatal brain from P0.5 to P7.5 (**Fig. S2**). Staining intensity was markedly lower at P0.5. At this stage, the lateral ventricle was surrounded by glycan-enriched white matter structures dorsally (corpus callosum, *cc*), posteriorly (fimbria of the hippocampus, *fi*), and ventrally (stria terminalis, *st*), but not anteriorly (**Fig. S2A**). By P7.5, all periventricular regions exhibited stronger and more robust staining (**Fig. S2B**). In contrast, no specific staining pattern was observed around the fourth ventricle. Given the supportive role of anionic glycans and proteoglycans, this spatially and temporally restricted development suggests a period of heightened susceptibility for the newborn brain ventricle to pathological insults.

Clinical trials commonly use dose escalation to evaluate AAV safety. Given the shared tropism of AAVs for the CSF-producing ChP, we administered two AAV doses into the neonatal brain ventricles in a common total of 1 μL volume to probe for adverse outcomes: 1 x 10^12^ GC/mL and 4 x 10^12^ GC/mL (**Fig. 2A**). Importantly, the doses we tested have both been reported in the literature for ICV injections of AAVs in preclinical models and are not generally considered excessive. First, we measured final brain wet weight at P7.5 (**Fig. 2B**) and monitored body weight over time (P2.5, P4.5, P7.5) (**Fig. 2C**). No significant alterations were observed in any treatment group, indicating general systemic and gross brain tolerance at this stage (**Fig. 2B, C**).

**Figure 2.**
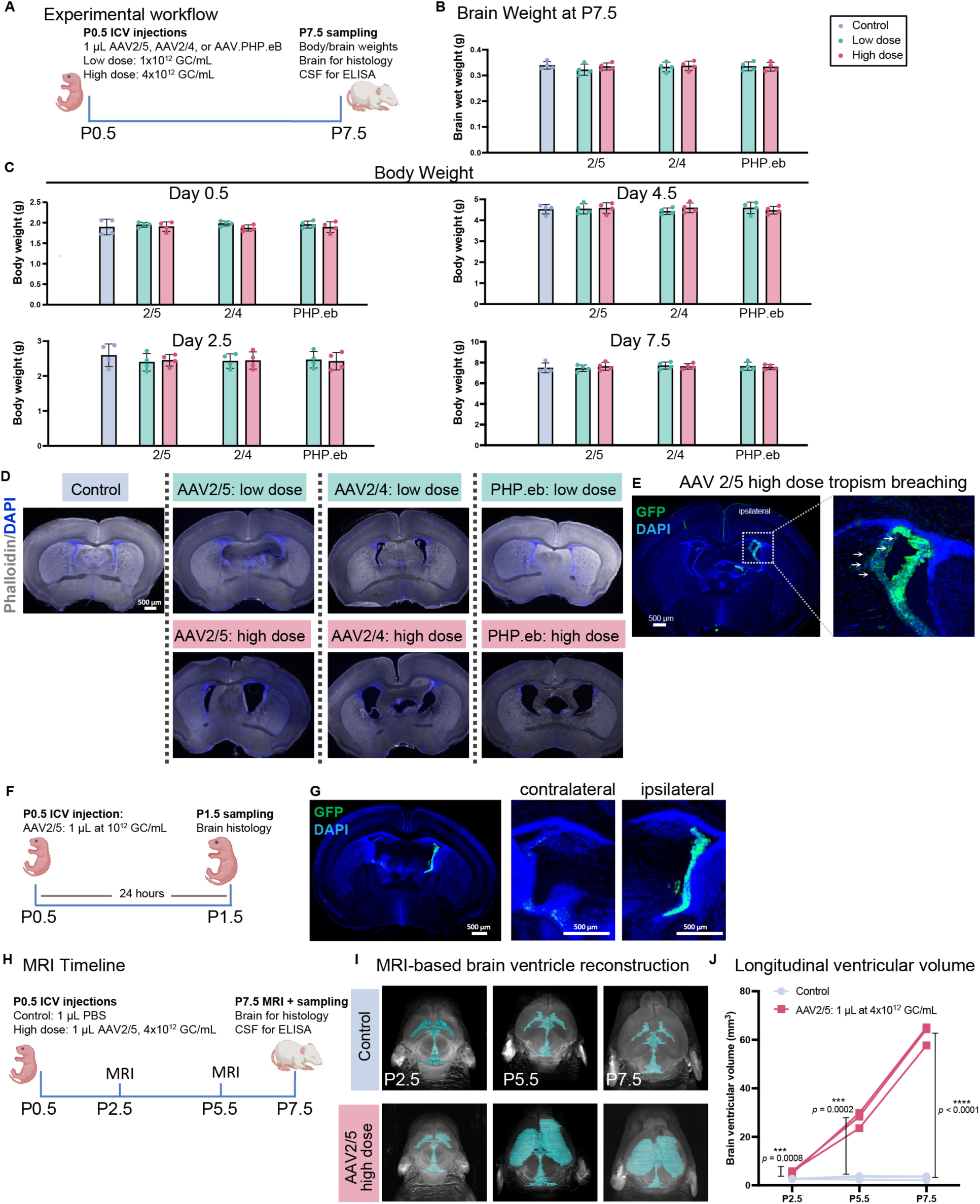
AAVs induce dose-dependent neonatal ventriculomegaly. **(A)** Experimental timeline to study the effects of ICV injection of AAV with escalating doses on neonatal brains. **(B)** Brain net weight measured on P7.5 post P0.5 ICV injections. **(C)** Body weight measured 2, 4, and 7 days post P0.5 ICV injections. **(D)** Representative coronal sections at the level of the posterior anterior commissure (pAC; dashed outline) showing lateral ventricle morphology in P7.5 brains following P0.5 ICV injections. **(E)** Representative image demonstrating the loss of cellular specificity ("permissive tropism") of high-dose AAV2/5, with ectopic GFP signal in ependymal regions. **(F, G)** Early AAV2/5 transfection kinetics. **(F)** Timeline for 24-hour analysis. **(G)** Representative image confirming robust ChP transfection 24 hours post-P0.5 injection (P1.5 brain). **(H-J)** Longitudinal MRI analysis of ventriculomegaly progression. **(H)** Timeline for serial scans following P0.5 ICV injections. **(I)** Representative 3D reconstructions of brain ventricles from control and high-dose AAV2/5-injected pups at 2, 5, and 7 days post-injection (P2.5, P5.5, P7.5). **(J)** Quantification of total ventricular volume across time points. (n = 3 per condition)

To screen for ventriculomegaly, we analyzed coronal brain sections at the level of the posterior anterior commissure (pAC), a standardized neuroanatomical landmark ventral to the anterior lateral ventricles (Ito et al. 2010; **Fig. 2D**). At low doses, AAVs caused no morphological alterations. In contrast, high-dose administration of all serotypes evidently enlarged the ventricles (**Fig. 2D**).

We further examined the cellular specificity of high-dose AAV2/5, which at a low dose showed exclusive tropism for CPECs (**Fig. 1D**). Histology revealed a loss of specificity, or "permissive" tropism, evidenced by new GFP signal in ependymal regions (**Fig. 2E**). Despite the higher dose, transfection remained largely ipsilateral, with minimal signal in the contralateral hemisphere. Overall, high-dose AAVs induced both dose-dependent neonatal ventriculomegaly and a permissive cellular tropism.

Considering the rapid postnatal development of brain environment (Pritschet et al. 2024; Ray et al. 2015), we asked whether the timing of injection influences AAV transfection pattern. Injecting the low dose at P1.5 instead of P0.5 increased transfection signal in the contralateral ChP (**Fig. S3A-B**). By P2.5 ICV injection resulted in nearly equal ChP transfection in both hemispheres, with no evident ipsilateral/contralateral difference (**Fig. S3C-D** ).

To characterize the progression of AAV-induced ventriculomegaly, we performed longitudinal MRI on pups receiving high doses of AAV2/5 on P0.5. We first confirmed robust AAV2/5 transfection within 24 hours of P0.5 injection (**Fig. 2F, G**). Serial scans were then performed 2, 5, and 7 days post-injection (P2.5, P5.5, P7.5) on pups administered a high-dose ICV injection on P0.5 (**Fig. 2H**). 3D reconstructions revealed severe enlargement of the lateral ventricles in AAV-injected brains compared to controls, with greater enlargement ipsilateral to the injection site (**Fig. 2I**), consistent with histology. Ventricular volume quantification confirmed the rapid progression of the condition, with significant enlargement detectable by day 2 that worsened through P7.5 (**Fig. 2J**).

To assess long-term health impacts, we tracked a cohort of P0.5 ICV-injected mice until adulthood (day 68). Only one mortality occurred (day 50, high-dose AAV2/5 group) (**Fig. S4A**). Body weight at 8 weeks post-injection showed no statistical significance overall, although an 18.5% reduction in body weight among male mice was found (*p* = 0.0967) (**Fig. S4B-C**).

### 3.3. AAVs induce dose-dependent CSF dyshomeostasis

Having established that AAV overdose induces ventriculomegaly, we next investigated the underlying mechanisms, focusing on two key metrics of brain ventricular health: neuroinflammation and ChP secretory function.

We first quantified the pro-inflammatory chemokine CCL2 (MCP-1) in CSF. ELISA revealed a dose-dependent increase in CSF CCL2 concentrations across all three AAV serotypes (**Fig. 3A**), with AAV.PHP.eB trending toward elevation (*p* = 0.175). Analysis of CSF transthyretin (TTR) revealed that high-dose AAV administration significantly decreased TTR concentrations irrespective of serotype, while low doses had no effect (**Fig. 3B**). These data suggest high-dose AAV disrupts CSF homeostasis by concurrently promoting inflammation and impairing ChP secretory function, whereas low doses are well-tolerated.

**Figure 3.**
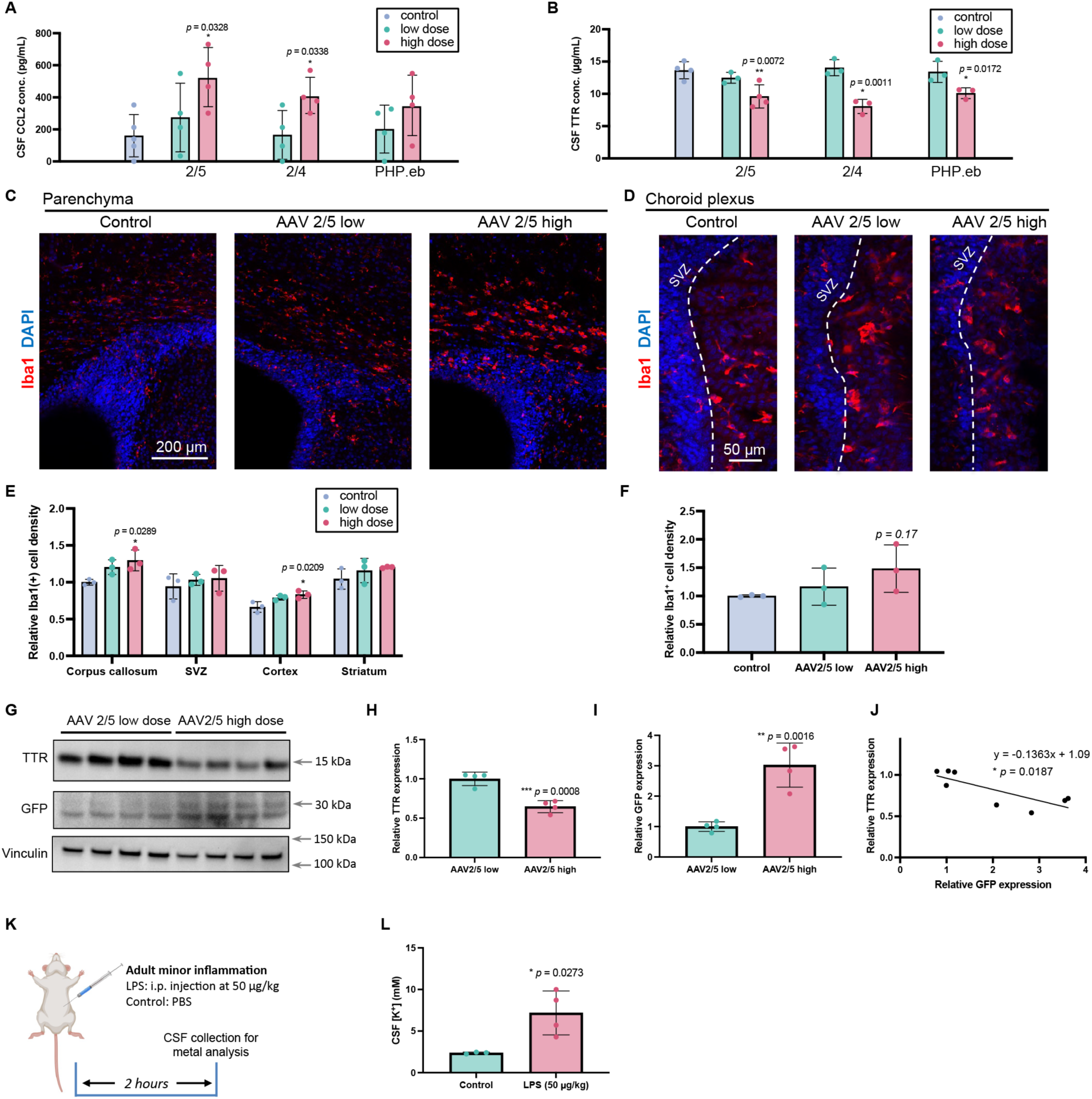
AAVs induce dose-dependent CSF dyshomeostasis. **(A)** CSF CCL2 concentrations measured by ELISA at P7.5 following P0.5 ICV injection of escalating AAV doses. **(B, C)** Parenchymal microglial response. **(B)** Representative Iba1 immunostaining in P7.5 brain sections. **(C)** Quantification of Iba1⁺ cell density in parenchymal regions. **(D, E)** ChP microglial response. **(D)** Representative Iba1 immunostaining in the ChP. **(E)** Quantification of Iba1⁺ cell density in the ChP. **(F)** CSF transthyretin (TTR) concentrations measured by ELISA at P7.5 following P0.5 ICV injection of escalating AAV doses. **(G–J)** ChP protein analysis by immunoblot. **(G)** Representative blot for TTR and GFP in ChP lysates. **(H)** Relative GFP expression. **(I)** Relative TTR expression. **(J)** Correlation analysis between GFP and TTR levels in individual samples. **(K, L)** CSF ion disruption following inflammation. **(K)** Experimental timeline for LPS-induced inflammation and CSF collection in adult mice. **(L)** CSF potassium (K⁺) concentration measured by ICP-MS.

Given the evidence of CSF inflammation, we assessed parenchymal microglial reactivity via Iba1 staining. Compared to controls, low-dose AAV2/5 did not change parenchymal Iba1, whereas high-dose administration increased microglial density in specific regions, i.e., the corpus callosum and cortex (**Fig. 3C, E**). Notably, despite being the primary site of AAV transfection, Iba1 density in the ChP itself was unaltered by either low or high AAV doses (**Fig. 3D, F**).

To further examine ChP dysfunction, we analyzed TTR protein levels in ChP tissue by immunoblotting following AAV2/5 administration. As expected, GFP expression was higher in the high-dose group (**Fig. 3G, I**). However, TTR levels in the ChP were significantly reduced in this group (**Fig. 3G, H**), consistent with CSF TTR levels (**Fig. 3B**). A correlational analysis confirmed a significant negative relationship between AAV transfection (evidenced by GFP levels) and TTR levels in individual samples (**Fig. 3J**; \**p* = 0.0187).

To integrate these two mechanisms, we investigated whether inflammation alters CSF ion concentrations because ventriculomegaly can be driven by osmotic dysfunction and CSF ion imbalance.

We induced minor systemic inflammation in adult mice via intraperitoneal LPS (50 µg/kg) (Osterhout et al. 2022) and collected CSF two hours later for metal analysis (**Fig. 3K**). ICP-MS revealed an approximate 3-fold increase in CSF potassium (K⁺) in LPS-treated mice (7.178 ± 2.635 mM) compared to controls (2.363 ± 0.1052 mM) (**Fig. 3L**).

Collectively, these results demonstrate that AAV overdose triggers concurrent inflammation in the CSF and parenchyma, disrupts critical ChP secretory function, and is associated with a rapid increase in CSF potassium during inflammation. Correcting this inflammatory cascade or the resulting ionic imbalance may offer a therapeutic strategy to mitigate AAV-induced ventriculomegaly.

### 3.4. AAV induces dose-dependent embryonic and adult ventriculomegaly

The pronounced ventriculomegaly observed in neonates after ICV delivery of high doses of AAVs led us to investigate whether embryonic brains at earlier developmental stages are also susceptible.

First, using a short timeline (**Fig. 4A**), we investigated whether AAV transgene expression is detectable shortly after in utero injection. Robust GFP was present in the embryonic ChP 24 hours after E13.5 administration (**Fig. 4B**).

**Figure 4.**
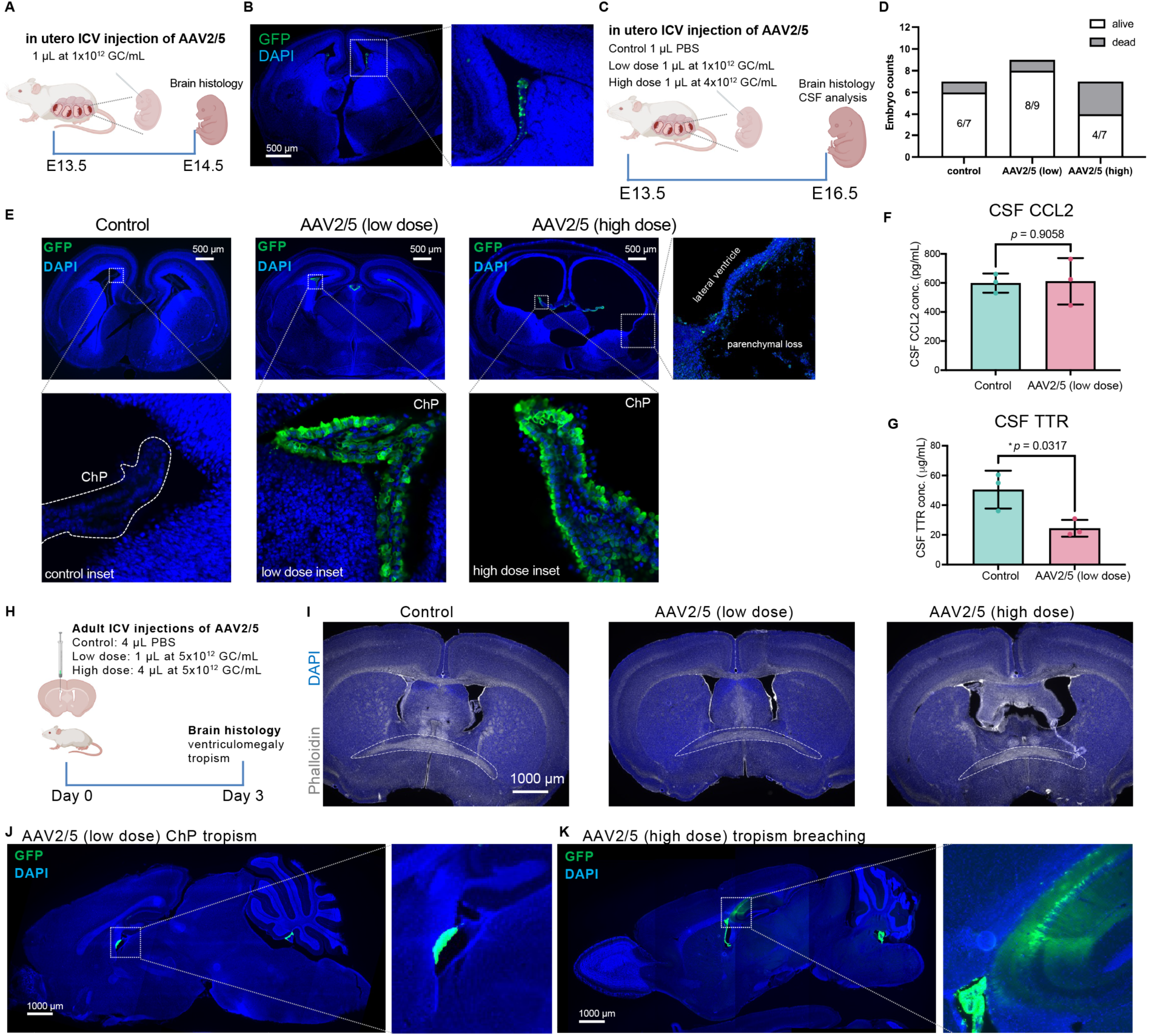
AAV induces dose-dependent embryonic and adult ventriculomegaly. **(A)** Experimental timeline for assessing acute AAV2/5 transfection 24 hours post in utero ICV injection. **(B)** Representative image confirming robust AAV2/5 transfection in the embryonic ChP at E14.5 (24 hours post-injection). **(C)** Timeline for evaluating the effects of escalating AAV2/5 doses via in utero ICV injection at E13.5, with analysis at E16.5. **(D)** Embryonic survival rates at E16.5. Numbers indicate the proportion of surviving embryos per litter. **(E)** Representative coronal sections of E16.5 embryonic brains showing transgene expression (GFP) and ventricular morphology. **(F, G)** CSF analysis from E16.5 embryos. **(F)** CCL2 concentrations. **(G)** Transthyretin (TTR) concentrations. **(H)** Timeline for assessing the effects of escalating AAV2/5 doses via ICV injection in adult mice (3-day endpoint). **(I)** Representative coronal sections at the level of the posterior anterior commissure (pAC; dashed outline) showing lateral ventricle morphology in adult brains. **(J, K)** Representative sagittal section demonstrating the differential cellular tropism of AAV2/5. **(J)** ChP-specific transfection by low-dose AAV2/5. **(K)** A more permissive tropism of high-dose AAV2/5, with ectopic GFP signal also observed in hippocampal CA1 region.

We then performed E13.5 in utero ICV injections of AAV2/5 at escalating doses and performed sampling three days later at E16.5 (**Fig. 4C**). High-dose AAV (1 μL at 4 x 10^12^ GC/mL) caused significant embryonic mortality (3 of 7 injected embryos; **Fig. 4D**). Surviving embryos in this group exhibited translucent, severely thinned brain parenchyma, precluding CSF collection, whereas control and low-dose (1 μL at 1 x 10^12^GC/mL ) embryos appeared largely normal. Histological analysis revealed that a low dose resulted in robust CPEC-specific tropism with normal ventricular morphology. In contrast, a high dose of ICV AAV2/5 induced ventriculomegaly, ectopic cellular tropism outside the ChP, and parenchymal loss (**Fig. 4E**). CSF analysis indicated that a low dose was well-tolerated with no increase in the inflammatory marker CCL2 in CSF (**Fig. 4F**) but did significantly reduce CSF transthyretin (TTR) concentration (**Fig. 4G**). These results collectively demonstrate a high vulnerability of the embryonic brain to AAV overdose.

Finally, we investigated whether AAV overdose induces ventriculomegaly in adult brains. We administered escalating doses of AAV2/5 via ICV injection and evaluated brains three days later (**Fig. 4H**). As expected, a low dose caused no ventricular alteration, whereas a high dose induced clear adult ventriculomegaly, though less severe compared to neonatal and embryonic stages (**Fig. 4I**). While low-dose AAV2/5 shows specific ChP tropism (**Fig. 4J**), the high-dose administration resulted in a permissive tropism, with robust ectopic transfection in hippocampal CA1 neurons (**Fig. 4K**).Together, these results show that embryonic brains are even more vulnerable to ICV delivery of AAVs at high doses.

## 4. Discussion

Data presented in this study demonstrate both common and distinct physiological and pathological properties of three CSF-relevant AAV serotypes following neonatal intracerebroventricular (ICV) injection. While AAV2/5 exhibited CPEC-specific tropism after low-dose administration, its administration at a high dose induced ventriculomegaly and more permissive tropism. In contrast, AAV2/4 demonstrated generally limited brain transfection, whereas ICV AAV.PHP.eB efficiently targeted both CPECs and ependymal cells. Notably, high-dose applications of AAV2/4 and AAV.PHP.eB also induced ventriculomegaly. Longitudinal MRI revealed the rapid progression of this condition post-injection, which was associated with CSF inflammation, microglial reactivity, and ChP dysfunction.

Furthermore, embryonic and adult brains were also susceptible to AAV-induced ventriculomegaly. Collectively, these findings underscore that although AAV-mediated gene therapy is promising for pediatric neurological disorders, its successful translation critically depends on meticulous serotype selection and precise dose titration to optimize efficacy and prevent iatrogenic injury.

AAV2/5 is established as a serotype with ChP tropism in embryonic and adult brains when delivered via ICV injections (Jang and Lehtinen 2022). Our histology data, demonstrating its selective transfection in the neonatal ChP, further corroborates this distinct targeting profile. Reported inconsistencies, such as studies observing parenchymal tropism following ICV administration, likely stem from critical methodological variations. For instance, while Chakrabarty et al. noted cortical transfection by AAV2/5, more prominent signals in the ChP were dismissed. Key experimental differences may explain these divergent findings, including a substantially higher viral dose (20-fold), surgical technique (freehand injection with a limited dorsal penetration that potentially results in a failure to reliably deliver virus into the CSF without delivering to the parenchyma), and mouse strain (B6C3F1/Tac vs. CD1). Importantly, when administered at an excessive dose in our study, AAV2/5 exhibited a loss of specificity, transfecting ependymal and hippocampal cells (**Fig. 2E and 4K**). This permissive tropism underscores a potential off-target risk, highlighting the necessity for precise dosing when considering this serotype for therapeutic applications. Our data further suggest that the state of the developing neonatal CSF/ventricular system critically influences transfection patterns. This is evidenced by a strongly ipsilateral signal from the P0.5 injection cohort that becomes increasingly more bilateral (also contralateral) by P1.5 and P2.5 injection cohorts (**Fig. S3**).

The earliest report of AAV2/4 described strong ependymal transfection in neonatal brains following ICV injection using an RSV promoter (Liu et al. 2005). However, subsequent studies from the same and other independent labs, primarily in adult mice, have reported robust transfection of both the ChP and ependyma by this serotype (Carrell et al. 2023; Dodge et al. 2010). These discrepancies in cellular tropism may be attributable to several experimental variables across studies. Notably, AAV2/4 constructs have utilized different promoters (e.g., RSV, VWA3A, or unspecified variants), which could significantly influence the observed reporter expression patterns. Combined with variations in host factors—such as mouse strain and age—these methodological differences suggest that the reported ependyma-specificity of AAV2/4 is not a fixed property. Therefore, establishing AAV2/4 as a reliably selective vector for ependymal targeting will require additional systematic investigation.

AAV.PHP.eB has been reported as highly effective for transfecting neonatal brain parenchyma, particularly neurons after intravenous or intraparenchymal delivery (Mathiesen et al. 2020). In our study using ICV delivery, however, this serotype showed robust transfection in the ChP and ependyma, with minimal parenchymal signals. Several methodological differences likely underlie this discrepancy: the timing of administration (P0.5 in our study vs. P1.5), a substantially higher viral dose (at least 10-fold greater), mouse strain (C57BL/6J vs. CD1), and potentially different ICV coordinates in prior reports.

Notably, strong ChP transduction—visible in representative images from prior studies (Chatterjee et al. 2022; Mathiesen et al. 2020)—has been largely unreported. These collective discrepancies highlight the necessity for a systematic, standardized evaluation of dose, injection parameters, viral source, and developmental timing in neonatal AAV studies.

The ChP directly interfaces with the CSF and is distinguished by a uniquely high density of brush border microvilli on its apical surface (Cornford et al. 1997; Spector et al. 2015). This structural specialization is directly evidenced by electron microscopy, which reveals a far greater microvillar density on ChP epithelial cells compared to other CSF-contacting cells, such as ependymal cells (Cornford et al. 1997; Spassky et al. 2005; Swiderski et al. 2012). Molecular analyses further support this unique architecture, demonstrated by the expression of microvilli-specific marker genes (e.g., Villin-1 and Villin-2) (Lein et al. 2007) and activity of the Villin promoter in the ChP (Rutlin et al. 2020).

Functionally, microvilli serve to vastly increase surface area and are known to facilitate endocytosis and the uptake of extracellular materials (Crawley et al. 2014). Consequently, this extensive microvillar network likely underpins the efficient transfection of the ChP observed for all intracerebroventricularly administered AAV serotypes in our study. Supporting this mechanism, our follow-up experiments using diameter-matched, negatively charged fluorescent microspheres—inert particles that mimic viral behavior—show exceptional accumulation in the ChP compared to other brain regions. Given the ChP’s inherent capacity for particulate uptake and its central role in CSF-ICF exchange, our findings highlight the critical importance of evaluating potential off-target effects in the ChP for any future therapeutic agent delivered directly into the CSF.

Pediatric hydrocephalus is a severe condition associated with significant short- and long-term clinical burdens, including neurological and cognitive impairment, social and educational obstacles, and increased mortality (Vinchon et al. 2012). While traditionally caused by genetic defects, infection, or cerebral hemorrhage (Kahle et al. 2024), hydrocephalus is now recognized as a critical potential side effect of therapeutics delivered directly to the CNS. For example, ICV delivery of an antisense oligonucleotide (ASO) targeting a KCNT1 mutation induced severe, progressive hydrocephalus with fatal outcomes in a pediatric cohort (Hayden 2022). Consistent with this clinical finding, our data demonstrate that AAV overdose—independent of serotype—induces neonatal ventriculomegaly, whereas lower doses appear safe. Longitudinal MRI revealed the rapid progression of this condition. Long-term follow-up into adulthood suggested further adverse outcomes, including mortality (one death on day 50) and an 18.5% reduction in body weight among male mice, although this latter difference was not statistically significant (*p* = 0.0967) (**Fig. S4**). With multiple clinical trials planning direct CSF delivery of AAVs to pediatric patients (e.g., NCT06948019, NCT06272149), our findings carry urgent relevance.

Given that AAV-induced ventriculomegaly also occurs in embryonic and adult brains, we underscore the indispensable need for precise viral dose titration and mandatory longitudinal monitoring of ventricular health via non-invasive imaging such as MRI.

CSF inflammation is implicated in hydrocephalus (Lolansen et al. 2021). Our data align with earlier reports and demonstrate dose-dependent neonatal CSF inflammation, as evidenced by elevated CSF CCL2 concentrations. Notably, this increase in CCL2 was not entirely dependent on viral transfection, as the minimally transducing serotype AAV2/4 also significantly raised CCL2 levels. Given the concurrent increase in parenchymal microglial density following high-dose AAV administration, a critical future investigation will be to test whether anti-inflammatory strategies can mitigate AAV-induced ventriculomegaly. Our findings also establish the AAV-overdose model as a valuable preclinical platform for screening potential hydrocephalus treatments. Furthermore, considering that even minor systemic inflammation elevated CSF potassium in adult mice, a promising therapeutic avenue may be to counteract this imbalance. Specifically, AAV2/5-mediated overexpression of NKCC1—which has shown ventricle-decreasing effects—could represent a targeted strategy to alleviate AAV-induced ventriculomegaly (Sadegh et al. 2023; Xu et al. 2021).

In summary, the data presented in this study elucidate both the utility and the potential safety risks of using AAV vectors for the pediatric population. Moving forward, our findings motivate two critical lines of investigation. First, brain alterations detected in neonates require longitudinal tracking further into later adulthood. This can be achieved through serial MRI to assess the permanence or reversibility of ventriculomegaly, complemented by behavioral assays to screen for lasting functional deficits. Second, the core mechanisms driving AAV-induced pathology remain to be defined. Key future experiments may include evaluating anti-inflammatory therapies and delineating disruptions in CSF dynamics to identify actionable therapeutic targets. Ultimately, our work reinforces the principle that AAV-mediated gene therapy must be advanced with a personalized medicine approach. Extrapolating findings across serotypes, doses, or species without rigorous validation carries significant risk and demands meticulous consideration.

## Acknowledgement

This work was supported by the Shurl and Kay Curci Foundation; the National Institutes of Health (R01 MH136258); and the Synthetic Neuroscience Grant Program of the Wu Tsai Neurosciences Institute at Stanford University (R.M.F) and the National Science Foundation (NSF) Graduate Research Fellowship Program (GRFP) and ARC Institute PhD Fellowship (B.Z.). Technical support was provided by the Stanford Gene Vector and Virus Core (RRID: SCR_023250). Longitudinal MRI was performed at the Stanford University Neurosciences Preclinical Imaging Laboratory (NPIL), supported by NIH grant S10OD025176, with assistance from Dr. Jin Hyung Lee. ICP-MS analysis was performed at Stanford University Environmental Measurements Facility with assistance from Dr. Guangchao Li. We also thank Dr. Phillip Beachy (Stanford University) for access to his fluorescence plate reader. Finally, we are grateful to all members of the Fame laboratory for their insightful discussions and proofreading.

## References

1. Alonso MI, Gato A, Moro JA, Barbosa E. 1998. Disruption of proteoglycans in neural tube fluid by β-D-xyloside alters brain enlargement in chick embryos. Anat Rec 252:499–508; doi:10.1002/(SICI)1097-0185(199812)252:4<499::AID-AR1>3.0.CO;2-1.

2. Amin S, Borrell V. 2020. The Extracellular Matrix in the Evolution of Cortical Development and Folding. Front Cell Dev Biol 8:604448; doi:10.3389/fcell.2020.604448.

3. Angata K, Huckaby V, Ranscht B, Terskikh A, Marth JD, Fukuda M. 2007. Polysialic Acid-Directed Migration and Differentiation of Neural Precursors Are Essential for Mouse Brain Development. Molecular and Cellular Biology 27:6659–6668; doi:10.1128/MCB.00205-07.

4. Carrell EM, Chen YH, Ranum PT, Coffin SL, Singh LN, Tecedor L, et al. 2023. VWA3A-derived ependyma promoter drives increased therapeutic protein secretion into the CSF. Molecular Therapy - Nucleic Acids 33:296–304; doi:10.1016/j.omtn.2023.07.016.

5. Chakrabarty P, Rosario A, Cruz P, Siemienski Z, Ceballos-Diaz C, Crosby K, et al. 2013. Capsid Serotype and Timing of Injection Determines AAV Transduction in the Neonatal Mice Brain. J. Qiu, ed PLoS ONE 8:e67680; doi:10.1371/journal.pone.0067680.

6. Chan KY, Jang MJ, Yoo BB, Greenbaum A, Ravi N, Wu W-L, et al. 2017. Engineered AAVs for efficient noninvasive gene delivery to the central and peripheral nervous systems. Nat Neurosci 20:1172–1179; doi:10.1038/nn.4593.

7. Chatterjee D, Marmion DJ, McBride JL, Manfredsson FP, Butler D, Messer A, et al. 2022. Enhanced CNS transduction from AAV.PHP.eB infusion into the cisterna magna of older adult rats compared to AAV9. Gene Ther 29:390–397; doi:10.1038/s41434-021-00244-y.

8. Cornford EM, Varesi JB, Hyman S, Damian RT, Raleigh MJ. 1997. Mitochondrial content of choroid plexus epithelium: Exp Brain Res 116:399–405; doi:10.1007/PL00005768.

9. Crawley SW, Mooseker MS, Tyska MJ. 2014. Shaping the intestinal brush border. Journal of Cell Biology 207:441–451; doi:10.1083/jcb.201407015.

10. Dodge JC, Treleaven CM, Fidler JA, Hester M, Haidet A, Handy C, et al. 2010. AAV4-mediated Expression of IGF-1 and VEGF Within Cellular Components of the Ventricular System Improves Survival Outcome in Familial ALS Mice. Molecular Therapy 18:2075–2084; doi:10.1038/mt.2010.206.

11. Flotte T, Carter B, Conrad C, Guggino W, Reynolds T, Rosenstein B, et al. 1996. A Phase I Study of an Adeno-Associated Virus-CFTR Gene Vector in Adult CF Patients with Mild Lung Disease. Johns Hopkins Children’s Center, Baltimore, Maryland. Human Gene Therapy 7:1145–1159; doi:10.1089/hum.1996.7.9-1145.

12. Hayden EC. 2022. Gene Treatment for Rare Epilepsy Causes Brain Side Effect in 2 Children. The New York Times, October 26.

13. Ito A, Shinmyo Y, Abe T, Oshima N, Tanaka H, Ohta K. 2010. Tsukushi is required for anterior commissure formation in mouse brain. Biochemical and Biophysical Research Communications 402:813–818; doi:10.1016/j.bbrc.2010.10.127.

14. Jang A, Lehtinen MK. 2022. Experimental approaches for manipulating choroid plexus epithelial cells. Fluids Barriers CNS 19:36; doi:10.1186/s12987-022-00330-2.

15. Kahle KT, Klinge PM, Koschnitzky JE, Kulkarni AV, MacAulay N, Robinson S, et al. 2024. Paediatric hydrocephalus. Nat Rev Dis Primers 10:35; doi:10.1038/s41572-024-00519-9.

16. Karimy JK, Reeves BC, Damisah E, Duy PQ, Antwi P, David W, et al. 2020. Inflammation in acquired hydrocephalus: pathogenic mechanisms and therapeutic targets. Nat Rev Neurol 16:285–296; doi:10.1038/s41582-020-0321-y.

17. Koyuncu OO, Hogue IB, Enquist LW. 2013. Virus Infections in the Nervous System. Cell Host & Microbe 13:379–393; doi:10.1016/j.chom.2013.03.010.

18. Lein ES, Hawrylycz MJ, Ao N, Ayres M, Bensinger A, Bernard A, et al. 2007. Genome-wide atlas of gene expression in the adult mouse brain. Nature 445:168–176; doi:10.1038/nature05453.

19. Liu G, Martins IH, Chiorini JA, Davidson BL. 2005. Adeno-associated virus type 4 (AAV4) targets ependyma and astrocytes in the subventricular zone and RMS. Gene Ther 12:1503–1508; doi:10.1038/sj.gt.3302554.

20. Lolansen SD, Rostgaard N, Oernbo EK, Juhler M, Simonsen AH, MacAulay N. 2021. Inflammatory Markers in Cerebrospinal Fluid from Patients with Hydrocephalus: A Systematic Literature Review. R. Rizzo, ed Disease Markers 2021:1–12; doi:10.1155/2021/8834822.

21. Mathiesen SN, Lock JL, Schoderboeck L, Abraham WC, Hughes SM. 2020. CNS Transduction Benefits of AAV-PHP.eB over AAV9 Are Dependent on Administration Route and Mouse Strain. Molecular Therapy - Methods & Clinical Development 19:447–458; doi:10.1016/j.omtm.2020.10.011.

22. Mendell JR, Al-Zaidy S, Shell R, Arnold WD, Rodino-Klapac LR, Prior TW, et al. 2017. Single-Dose Gene-Replacement Therapy for Spinal Muscular Atrophy. N Engl J Med 377:1713–1722; doi:10.1056/NEJMoa1706198.

23. Osterhout JA, Kapoor V, Eichhorn SW, Vaughn E, Moore JD, Liu D, et al. 2022. A preoptic neuronal population controls fever and appetite during sickness. Nature 606:937–944; doi:10.1038/s41586-022-04793-z.

24. Paulson JN, Williams BL, Hehnly C, Mishra N, Sinnar SA, Zhang L, et al. 2020. *Paenibacillus* infection with frequent viral coinfection contributes to postinfectious hydrocephalus in Ugandan infants. Sci Transl Med 12:eaba0565; doi:10.1126/scitranslmed.aba0565.

25. Pritschet L, Taylor CM, Cossio D, Faskowitz J, Santander T, Handwerker DA, et al. 2024. Neuroanatomical changes observed over the course of a human pregnancy. Nat Neurosci 27:2253–2260; doi:10.1038/s41593-024-01741-0.

26. Ray S, Tzeng R-Y, DiCarlo LM, Bundy JL, Vied C, Tyson G, et al. 2015. An Examination of Dynamic Gene Expression Changes in the Mouse Brain During Pregnancy and the Postpartum Period. G3 (Bethesda) 6:221–233; doi:10.1534/g3.115.020982.

27. Rutlin M, Rastelli D, Kuo WT, Estep JA, Louis A, Riccomagno MM, et al. 2020. The Villin1 Gene Promoter Drives Cre Recombinase Expression in Extraintestinal Tissues. Cellular and Molecular Gastroenterology and Hepatology 10:864–867.e5; doi:10.1016/j.jcmgh.2020.05.009.

28. Sadegh C, Xu H, Sutin J, Fatou B, Gupta S, Pragana A, et al. 2023. Choroid plexus-targeted NKCC1 overexpression to treat post-hemorrhagic hydrocephalus. Neuron 111:1591–1608.e4; doi:10.1016/j.neuron.2023.02.020.

29. Schnaar RL, Gerardy-Schahn R, Hildebrandt H. 2014. Sialic Acids in the Brain: Gangliosides and Polysialic Acid in Nervous System Development, Stability, Disease, and Regeneration. Physiological Reviews 94:461–518; doi:10.1152/physrev.00033.2013.

30. Spassky N, Merkle FT, Flames N, Tramontin AD, García-Verdugo JM, Alvarez-Buylla A. 2005. Adult Ependymal Cells Are Postmitotic and Are Derived from Radial Glial Cells during Embryogenesis. J Neurosci 25:10–18; doi:10.1523/JNEUROSCI.1108-04.2005.

31. Spector R, Keep RF, Robert Snodgrass S, Smith QR, Johanson CE. 2015. A balanced view of choroid plexus structure and function: Focus on adult humans. Experimental Neurology 267:78–86; doi:10.1016/j.expneurol.2015.02.032.

32. Swiderski RE, Agassandian K, Ross JL, Bugge K, Cassell MD, Yeaman C. 2012. Structural defects in cilia of the choroid plexus, subfornical organ and ventricular ependyma are associated with ventriculomegaly. Fluids Barriers CNS 9:22; doi:10.1186/2045-8118-9-22.

33. Vinchon M, Rekate H, Kulkarni AV. 2012. Pediatric hydrocephalus outcomes: a review. Fluids Barriers CNS 9:18; doi:10.1186/2045-8118-9-18.

34. Weinhold B, Seidenfaden R, Röckle I, Mühlenhoff M, Schertzinger F, Conzelmann S, et al. 2005. Genetic Ablation of Polysialic Acid Causes Severe Neurodevelopmental Defects Rescued by Deletion of the Neural Cell Adhesion Molecule. Journal of Biological Chemistry 280:42971–42977; doi:10.1074/jbc.M511097200.

35. Wright JF. 2008. Manufacturing and characterizing AAV-based vectors for use in clinical studies. Gene Ther 15:840–848; doi:10.1038/gt.2008.65.

36. Xu H, Fame RM, Sadegh C, Sutin J, Naranjo C, Della Syau, et al. 2021. Choroid plexus NKCC1 mediates cerebrospinal fluid clearance during mouse early postnatal development. Nat Commun 12:447; doi:10.1038/s41467-020-20666-3.

