## Supplementary figures 1-5 for "Adeno-associated viruses (AAVs) induce dose-dependent neonatal ventriculomegaly following intracerebroventricular administration"

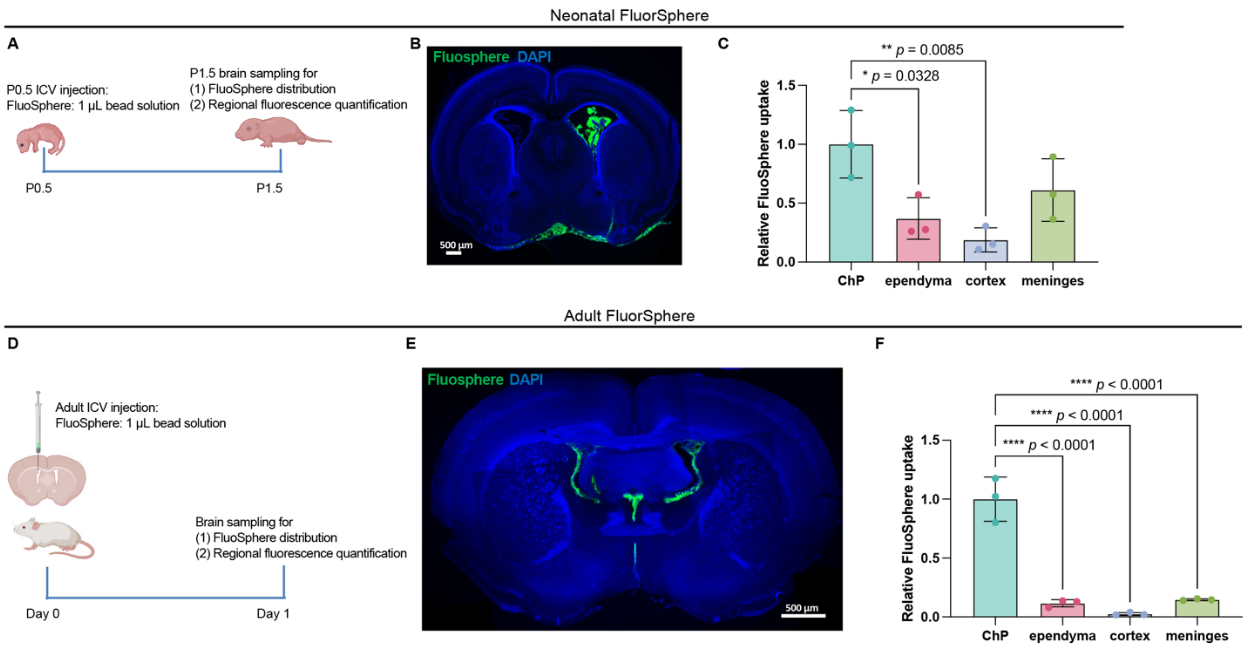


**Supplementary Figure 1. Cellular uptake of AAV size- and charge-matched particles following ICV administration: the choroid plexus as the predominant site.**

**(A)** Experimental timeline for ICV injection of FluoSphere beads in neonatal (P0.5) pups.

**(B)** Representative image showing the distribution of FluoSpheres (green) 24 hours post-ICV injection in a P1.5 neonatal brain.

**(C)** Quantitative analysis of FluoSphere fluorescence intensity across brain regions 24 hours post-injection in neonates (n = 3).

**(D)** Experimental timeline for ICV injection of FluoSphere beads in adult mice.

**(E)** Representative image showing FluoSphere distribution (green) 24 hours post-ICV injection in adult mouse brain.

**(F)** Quantitative analysis of FluoSphere fluorescence intensity across brain regions in adult mouse brain 24 hours post-injection (n = 3).

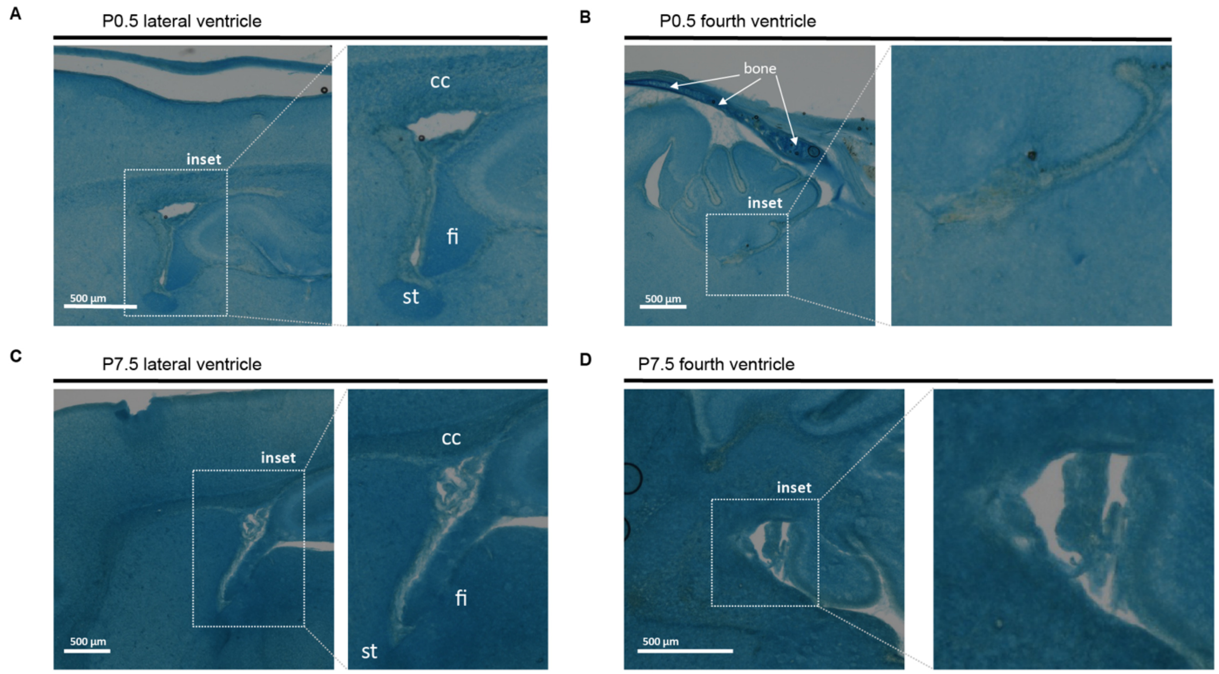


**Supplementary Figure 2. Neonatal development of brain extracellular matrix anionic glycan molecules by Alcian blue staining.**

**(A)** Alcian blue staining of P0.5 sagittal brain sections for the lateral and fourth ventricles. Insets highlight the characteristic periventricular staining pattern. *st*: stria terminalis; *fi*: fimbria; *cc*: corpus callosum.

**(B)** Alcian blue staining of P7.5 sagittal brain sections, showing increased staining intensity in periventricular regions. *st*: stria terminalis; *fi*: fimbria; *cc*: corpus callosum.

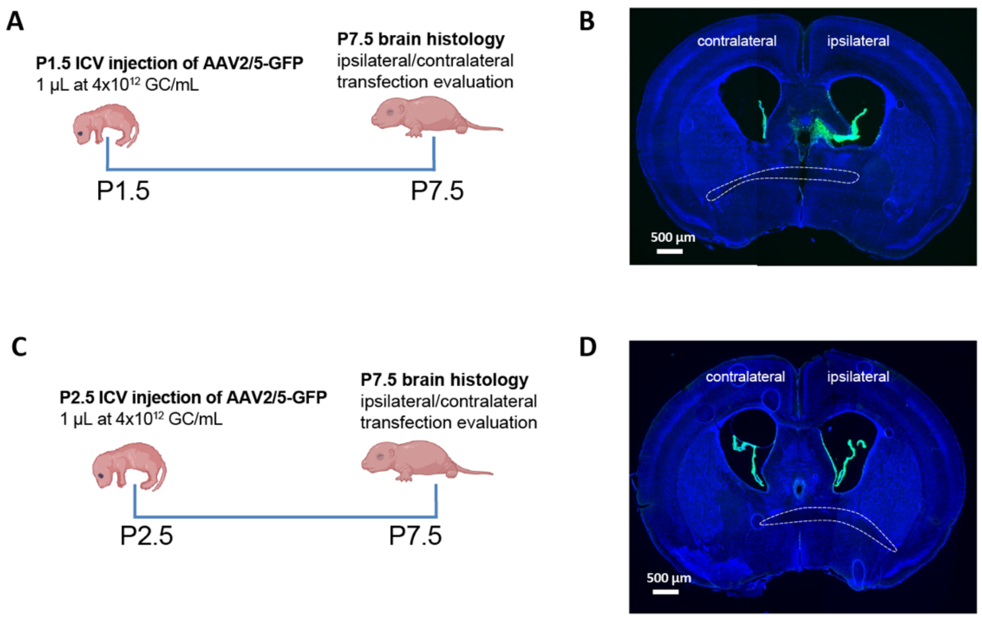


**Supplementary Figure 3. Neonatal ICV injection timing influences the ipsilateral/contralateral transfection pattern, but not the ventriculomegaly outcomes.**

**(A)** Experimental timeline for P1.5 ICV injection of high-dose AAV2/5 and analysis at P7.5.

**(B)** AAV2/5 transfection pattern (GFP) in P7.5 brains. The dashed line marks posterior anterior commissure (pAC).

**(C)** Experimental timeline for P2.5 ICV injection of high-dose AAV2/5 and analysis at P7.5.

**(D)** AAV2/5 transfection pattern (GFP) in P7.5 brains. The dashed line marks the pAC.

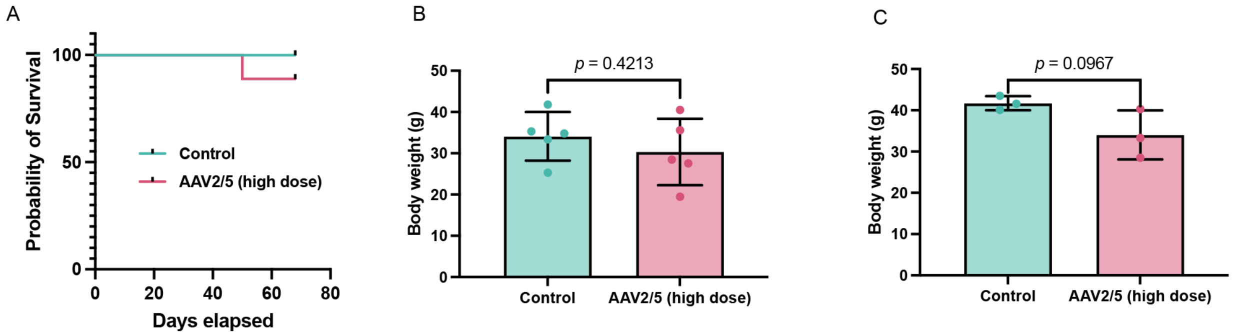


**Supplementary Figure 4. Long-term health assessment following neonatal ICV injection.**

**(A)** Survival curve tracked to postnatal day 68 following P0.5 ICV injections of high-dose AAV2/5 (n = 7-8 per condition).

**(B, C)** Body weight of female (n = 5 per condition) **(B)** and male (n = 3 per condition) **(C)** mice measured 8 weeks post-P0.5 ICV injection.

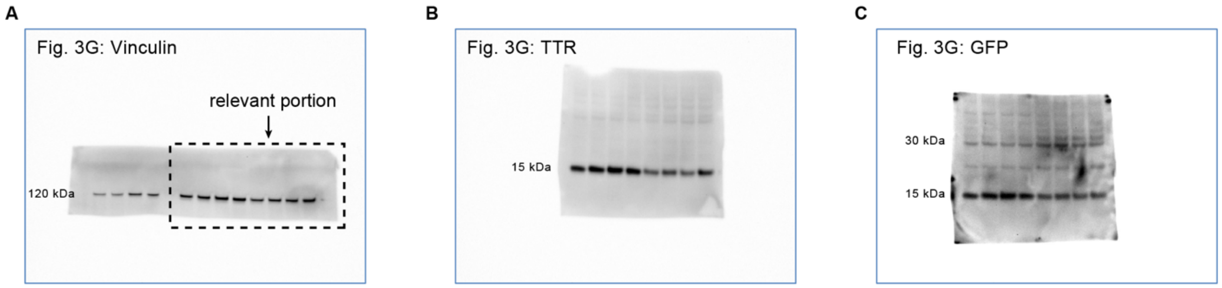


**Supplementary Figure 5. Full western blot membranes corresponding to Figure 3G.**

**(A)** Vinculin immunoblot.

**(B)** Transthyretin (TTR) immunoblot.

**(C)** The membrane from (B) was stripped and re-probed for GFP.
